# ChromeBat: A Bio-Inspired Approach to 3D Genome Reconstruction

**DOI:** 10.1101/2021.03.04.433995

**Authors:** Brandon Collins, Philip N. Brown, Oluwatosin Oluwadare

## Abstract

**Background:** With the advent of Next Generation Sequencing and the Hi-C experiment, high quality genome-wide contact data is becoming increasingly available. This data represents an empirical measure of how a genome interacts inside the nucleus. Genome conformation is of particular interest as it has been experimentally shown to be a driving force for many genomic functions from regulation to transcription. Thus, the Three Dimensional Genome Reconstruction Problem seeks to take Hi-C data and produce the complete physical genome structure as it appears in the nucleus for genomic analysis.

**Results:** We propose and develop a novel method to solve the Chromosome and Genome Reconstruction problem based on the Bat Algorithm which we called ChromeBat. We demonstrate on real Hi-C data that ChromeBat is capable of state of the art performance. Additionally, the domain of Genome Reconstruction has been criticized for lacking algorithmic diversity, and the bio-inspired nature of ChromeBat contributes algorithmic diversity to the problem domain.

**Conclusions:** ChromeBat is an effective approach at solving the Genome Reconstruction Problem. The source code and usage guide can be found here: https://github.com/OluwadareLab/ChromeBat.

## 1 Background

### 1.1 The Conformation Capture Assays

As DNA sequencing technology matures, so have questions surrounding how gene expression is functionally accomplished. It is well understood that genes require their associated regulators to function properly. However, DNA sequencing shows that a gene’s regulators may be many base pairs (bp) from the gene it regulates [1]. One experimentally proven mechanism to account for this disparity is the three-dimensional (3D) structure of the genome [2]. In particular, a gene’s regulator may be far in terms of linear base pairs, but in 3D space could be quite local. Thus, it is imperative to understand a genome’s structure in 3D space as it is a mechanism for gene function.

There is a rich history of assay development to understand 3D genomic structure. Recently, the rise of genome interaction measurement techniques based on a “all versus all read-pair interaction profiling”[3, 4], have enabled algorithmic approaches to reconstruct the genome. The first of these techniques, known as Hi-C [5], is summarized as follows: crosslink the chromatins using a fixative agent, digest the chromatin with a 4 or a 6 base cutter restrictive enzyme, apply biotin labels at the ends of the chromatins, relitigate the chromatins in dilute conditions, purify and shear DNA, and perform biotin pull-down [6]. Next, Next Generation Sequencing (NGS) technology is used for paired-end sequencing. The resulting reads are mapped to a reference genome, and filtered. This step results in the creation of an Interaction Frequency (IF) matrix, or contact matrix, representing relative levels of closeness of different portions of DNA called loci or bins. The length of the bins is called the resolution of the contact matrix. Hence, a bin with 1,000,000 base pairs has a resolution of 1mb. The Hi-C method’s main innovation is that it can supply data across the entire genome, allowing for 3D reconstructions at both the chromosome and genome-wide levels. Hi-C and related techniques are limited only by read depth and resolution restrictions presented by current sequencing technology. However, as NGS techniques steadily improve both in cost efficiency and throughput, Hi-C is poised to deliver genome-wide interaction datasets with ever increasing resolutions for bioinformatic analysis.

### 1.2 The Genome Reconstruction Problem

In this work we focus on using Hi-C data to solve the 3D genome reconstruction problem (3D-GRP). Formalized in [7], the 3D-GRP problem is defined as follows. First a Hi-C experiment is performed and a *contact matrix* is produced. A contact matrix is a square symmetric *n* × *n* matrix, where *n* is the number of loci at a given resolution. A solution to the 3D-GRP is set a (*x, y, z*) coordinates, one for each loci. A good solution will conform to the contact matrix from the Hi-C data. Approaches to solve this problem can fit into one of three categories. These are distance-based approaches, contact based approaches, and probabilistic approaches, which we briefly survey here [8].

Distance-based approaches feature two steps: first the contact matrix must be converted to a *distance matrix*, and then an optimization technique is applied. Focusing on the first step, the contact matrix is converted to a distance matrix via an inverse relationship based on a constant *α*, called the conversion factor, typically in the range (0, 3] [9]. Early approaches assumed an inverse relationship between distance, such as the 5C3D method developed in [10]. However, [11] demonstrated that the relationship between interaction frequency and distance can vary between experimental procedures and organisms illuminating the need for a principled method for picking *α*. One proposed solution [12] is to use microscopy data from FISH as a ground truth to assist in the interaction frequency to distance conversion process. Another approach [13] is to use a search algorithm to select a suitable *α* for each experiment.

Once the contact matrix has been converted to a distance matrix, the distance approaches proceed with optimization. One of the most popular choices [9] is to use a multidimensional scaling (MDS) approach [14]. This is the approach used in the classical 5C3D technique as well as more modern approaches such as miniMDS [15]. Another promising optimization process showcased in 3Dmax [16] involves formulating the problems in terms of maximum likelihood and solving it using an iterative technique such as gradient ascent. Other distance based methods include HSA [17], ChromeSDE [11], Shrec3D [18], Chromosome3D [19], and LorDG [20]. The second class of techniques are known as contact-based approaches. Unlike distance-based approaches, contact approaches derive a 3D structure directly from e Hi-C contact matrix. This is inherently advantageous as no assumption about a distance interaction frequency relationship needs to be made. The most straightforward of these approaches is known as MOGEN [21] which directly applies the gradient ascent optimization technique seeking to satisfy interaction thresholds given by the data. Purported to be robust against noise [8], it should be noted that noise and experimentally induced biases are highly nontrivial to handle. To mitigate this, contact based approaches have incorporated other sources of data such as fluorescence in situ hybridization (FISH) [22] as well as Lamina-associated Domains (LADs) [23].

The final class of genome reconstruction techniques is known as probability based methods. These methods function by defining a probability measure for contact frequencies. A major advantage for these methods is that uncertainty and bias in Hi-C data can be handled natively by a probabilistic method. Typically, probability-based approaches are ensemble techniques [8]. This entails the method will output a population of models whose average is representative of the Hi-C data, which intuitively makes sense as Hi-C data is usually an average of many cells. The classical probabilistic method is known as MCMC5C [24] which generates an ensemble of models based on Markov Chain Monte Carlo sampling. Another example of a probability based approach is PASTIS [25].

Although these techniques vary greatly in performance, computational efficiency, and output file format, they all represent a solution to the 3D-GRP problem. Unfortunately, validating these solutions has itself proven to be challenging. For example, consider using a norm that measures the distance between the distance matrix and a proposed structure’s induced distance matrix to evaluate these solutions. Immediately, we must assume that some *α* exists such that the contact matrix can be converted to ground truth distances between all loci and that our solution finds it. Additionally, it is plausible that a more complex formula is better at converting interaction frequency to distance. Secondly, it would only be valid on single cell Hi-C data or else the objective function would seek an average of genomic structures that does not exist. In practice, the 3D-GRP solutions are validated using known genomic structures, other data such as FISH [8], or a simulated dataset [26] where the ground truth structure is known. Thus, regardless of the quality of solution presented for the 3D-GRP, validating that the procedure is generating genomic structures representative of actual cells remains an open question.

### 1.3 Motivating ChromeBat

The 3D-GRP has become an important problem in genomics and computational biology due to the structural impact on genomic function. In this work we propose and explore a Bat Algorithm [27] based approach. The motivation for this approach is two-fold. First, existing approaches have been criticized for lacking algorithmic diversity [9]. Broadly speaking, the Bat Algorithm is a metaheuristic optimization algorithm, which is a family of algorithms that has seen some application to the problem. Notable applications of metaheuristic algorithms to the 3D-GRP are simulated annealing (SA) [28, 19] and genetic algorithms [29]. Specifically, the Bat Algorithm is a nature-based swarming algorithm which has seen no application on this problem, thus addressing the complaint of poor algorithmic diversity.

The second motivation is that the 3D-GRP is a high dimensional optimization problem. For example, consider the human genome reconstruction problem at relatively low resolution such as 1 megabase (mb). The human genome has approximately 3,200,000,000 nucleotides; at 1mb resolution this results in about 3,200 loci. Because the problem calls for selecting an *x, y, z* coordinate for each, there will be about 9,600 parameters to optimize, making it a high dimensional problem. Critically, increasing the resolution dramatically increases the dimensionality of the optimization problem. Because the Bat Algorithm has successfully been applied in other high dimensional settings [30], we propose it for the 3D-GRP.

## 2 Results

### 2.1 Comparison with Metaheuristic Methods

In the literature we have found six metaheuristic methods to compare our method against. These are: Gen3D [29], PGS [31], Chrom3D [23], 3D-GNOME [32], Chromosome3D [19], and HSA [17]. Of these, three are distance based (3D-GNOME, Chromosome3D, HSA), two are contact based (Gen3D, Chrom3D), and one is probability based (PGS).

Among these methods, Gen3D utilizes a genetic algorithm approach while the rest of the approaches are based on Simulated Annealing (SA). Unfortunately, PGS, Chrome3D, gen3D and 3D-GNOME all require more input data in addition to the contact matrices, so we could not compare against them. Regardless, we compare ChromeBat on GM06990 and GM12878 cell lines as discussed in Section 5.5 with HSA and Chromosome3D.

In the case of the GM06990 cell line, seen in Figure 1, ChromeBat has the highest SCC across all chromosomes evaluated. In particular, ChromeBat outperforms all other metaheuristic methods by at least 5% in chromosomes 14-18. For the GM12878 cell line, with results in Figure 2, we find that Chromebat is the most performant method in terms of SCC on the higher chromosomes while Chromosome3D produces the best structures on the remaining chromosomes.

**Figure 1.**
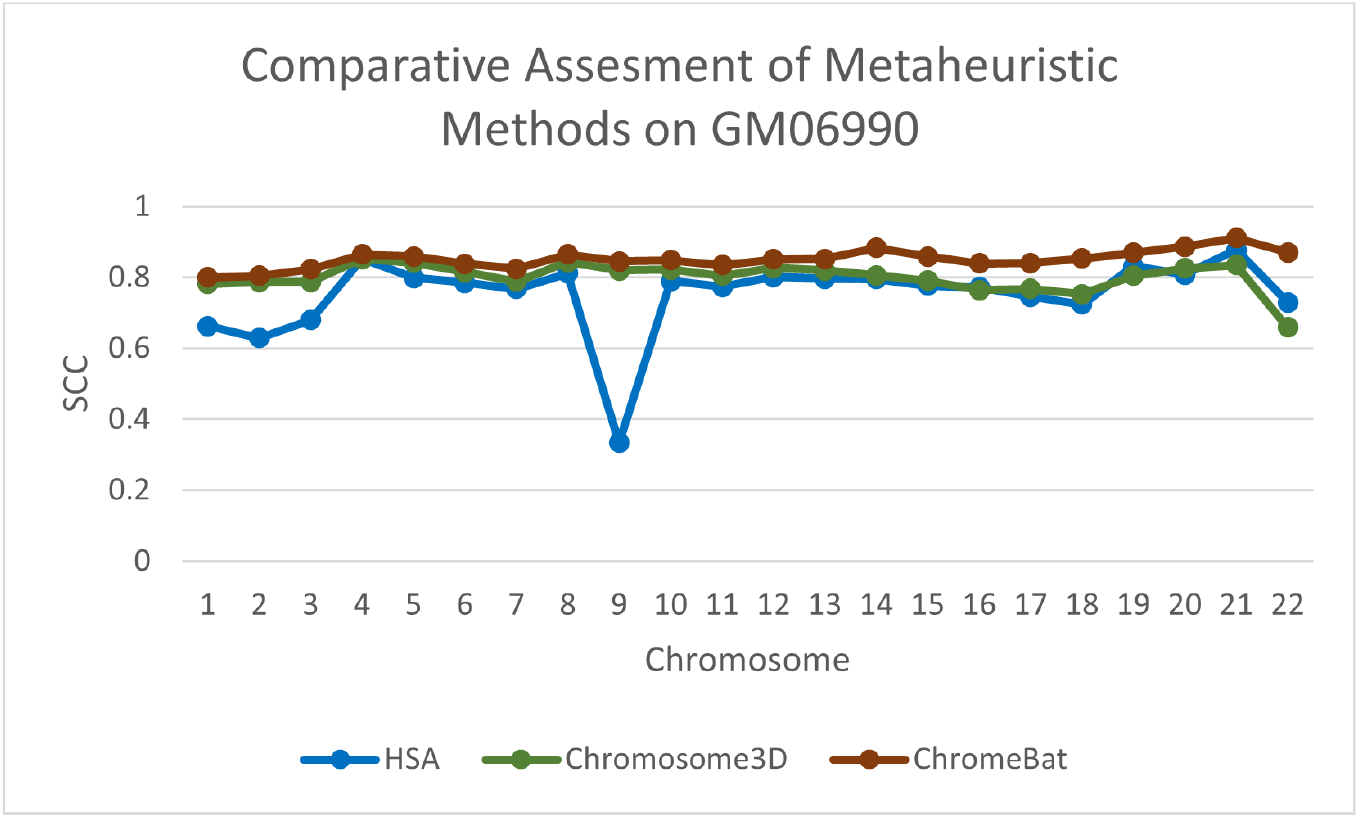
Comparative Assessment of Metaheuristic Methods on GM06990. A comparison of the Spearman Correlation Coefficient between metaheuristic methods ChromeBat, HSA, and Chromosome3D. This experiment is done on the first 22 chromosomes of the GM06990 cell line at 1mb resolution.

**Figure 2.**
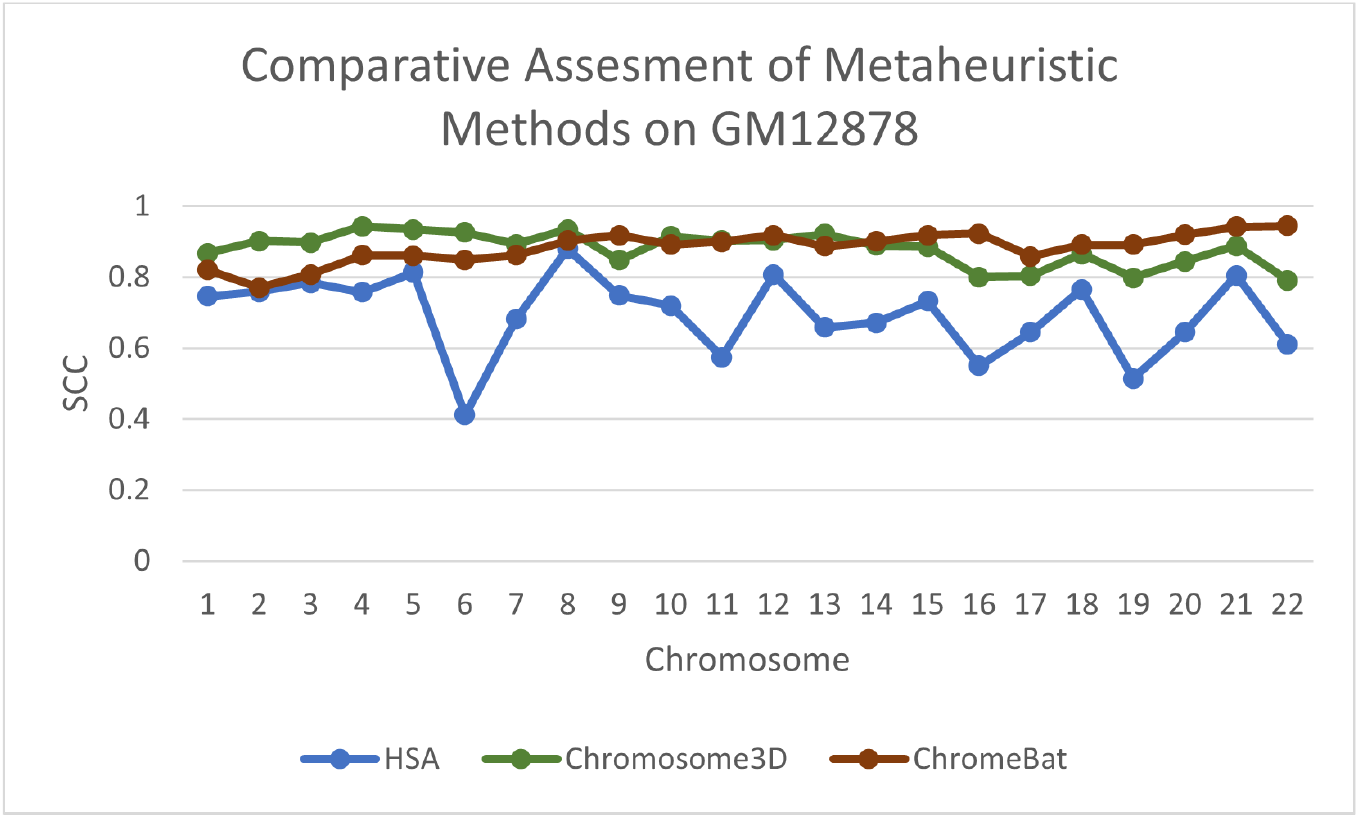
Comparative Assessment of Metaheuristic Methods on GM12878. A comparision of the Spearman Correlation Coefficient (SCC) between metaheuristic methods ChromeBat, HSA and Chromosome3D. This experiment is done on the first 22 chromosomes of the GM12878 cell line at 1mb resolution.

### 2.2 Comparison with Existing 3D-GRP Methods in Literature

To verify that ChromeBat is not only competitive among metaheuristic methods, we compare against eight literature methods on the GM06990 and GM12878 cell lines previously discussed. The methods we compare against are 3Dmax [16], HSA [17], ChromSDE [11], Pastis [25], ShRec3D [18], Chromosome3D [19], LorDG [20], and MOGEN [21]. For this comparison we take advantage these published results and computed the SCC metric scores for the remaining methods.

The results on the GM06990 cell line can be seen in Figure 3 or [see Additional file 6]. Overall, ChromeBat performs competitively across the board within a close margin of 3Dmax and ChromSDE on every chromosome. Of particular interest is ChromeBat’s performance on chromosome 20 where it has more then 3% performance improvement compared to the next best method. Secondly, we compare on the GM12878 cell line whose results may be seen in Figure 4 or [see Additional file 7]. Interestingly, all methods across the board score higher SCC on the GM12878 cell line relative to the GM06990 cell line. ChromeBat remains competitive although the performance suffers in the earlier chromosomes. Note that ChromeBat performs comparatively better on the GM06990 cell line that is more difficult to reconstruct across all methods.

**Figure 3.**
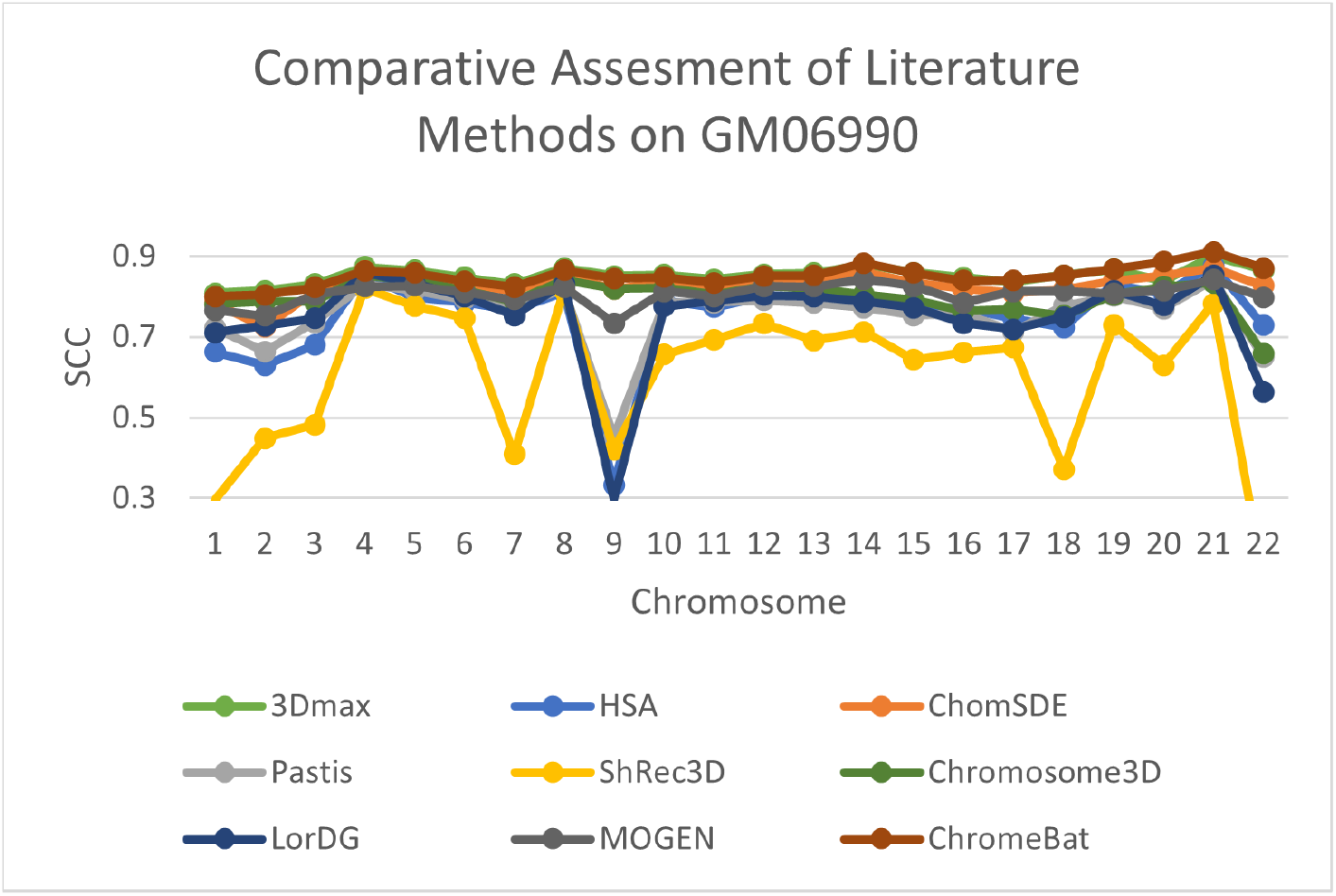
Comparative Assessment of Literature Methods on GM06990. A comparison of the Spearman Correlation Coefficient between a selection of literature methods. This experiment is done on the first 22 chromosomes of the GM06990 cell line at 1mb resolution.

**Figure 4.**
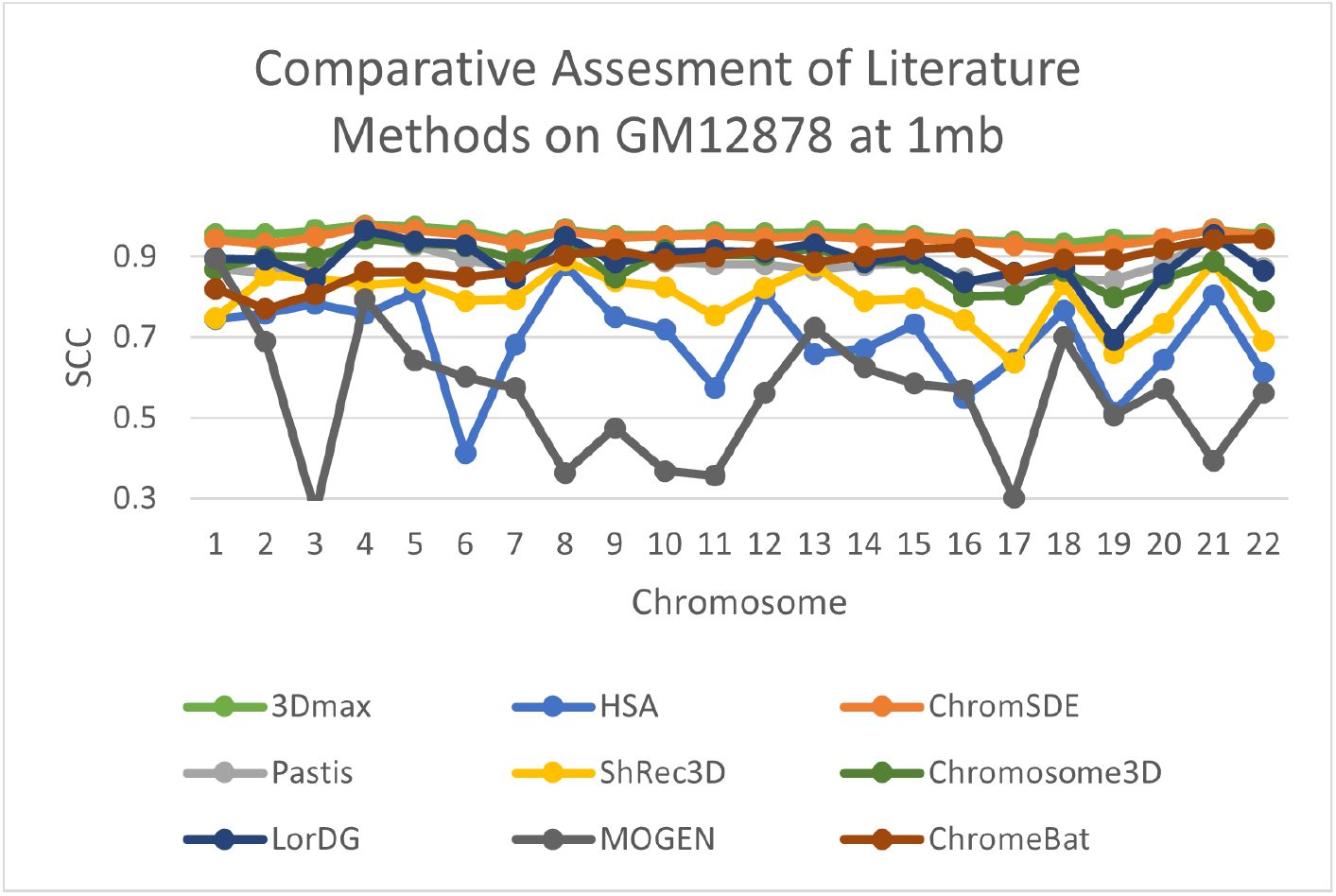
Comparative Assessment of Literature Methods on GM12878. A comparison of the Spearman Correlation Coefficient between a selection of literature methods. This experiment is done on the first 22 chromosomes of the GM12878 cell line at 1mb resolution.

### 2.3 Consistency

In Section 5.3 we note that the method appears to struggle with consistency under certain parameters. In this section we investigate how parameter choice interacts with the consistency of ChromeBat. We perform two experiments on the GM12878 cell line using the parameters specified in Table 1 with the exception that we take `structs = 20` and fix *p* = 0.002 in the first experiment and *p* = 0.004 in the second. The results are shown in Figures 5 and 6. Notice on the lower chromosomes with *p* = 0.002 ChromeBat displays high variability in performance between runs with the same parameters. However, simultaneously *p* = 0.002 produces more consistent and better structures on the later chromosomes than the *p* = 0.004 experiment, despite its poor performance on chromosomes 1-6. This reinforces the need to search across both *α* and *p*. Further, it shows that when the parameters of ChromeBat are well tuned, it can produce consistent and performant structures. It is then also important to generate multiple structures with the same parameters after the search is done, as the consistency of the produced structures reveals how well the hyperparameters are suited to the particular problem instance at hand.

**Table 1.**
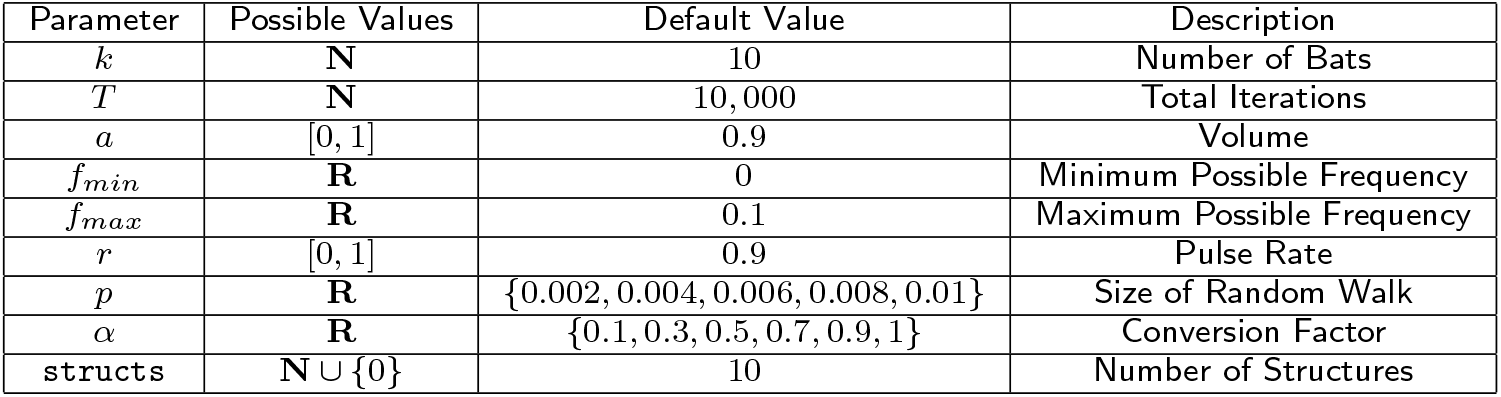
Hyperparameters in ChromeBat. Note in the original work certain variables such as *r, a* were proposed to change as the algorithm progressed. In ChromeBat they are constant throughout. For more information about the default value selection see Section 5.3.

| Parameter | Possible Values | Default Value | Description |
| --- | --- | --- | --- |
| $k$ | $\mathbf{N}$ | 10 | Number of Bats |
| $T$ | $\mathbf{N}$ | 10,000 | Total Iterations |
| $a$ | $[0, 1]$ | 0.9 | Volume |
| $f_{min}$ | $\mathbf{R}$ | 0 | Minimum Possible Frequency |
| $f_{max}$ | $\mathbf{R}$ | 0.1 | Maximum Possible Frequency |
| $r$ | $[0, 1]$ | 0.9 | Pulse Rate |
| $p$ | $\mathbf{R}$ | $\{0.002, 0.004, 0.006, 0.008, 0.01\}$ | Size of Random Walk |
| $\alpha$ | $\mathbf{R}$ | $\{0.1, 0.3, 0.5, 0.7, 0.9, 1\}$ | Conversion Factor |
| structs | $\mathbf{N} \cup \{0\}$ | 10 | Number of Structures |

**Figure 5.**
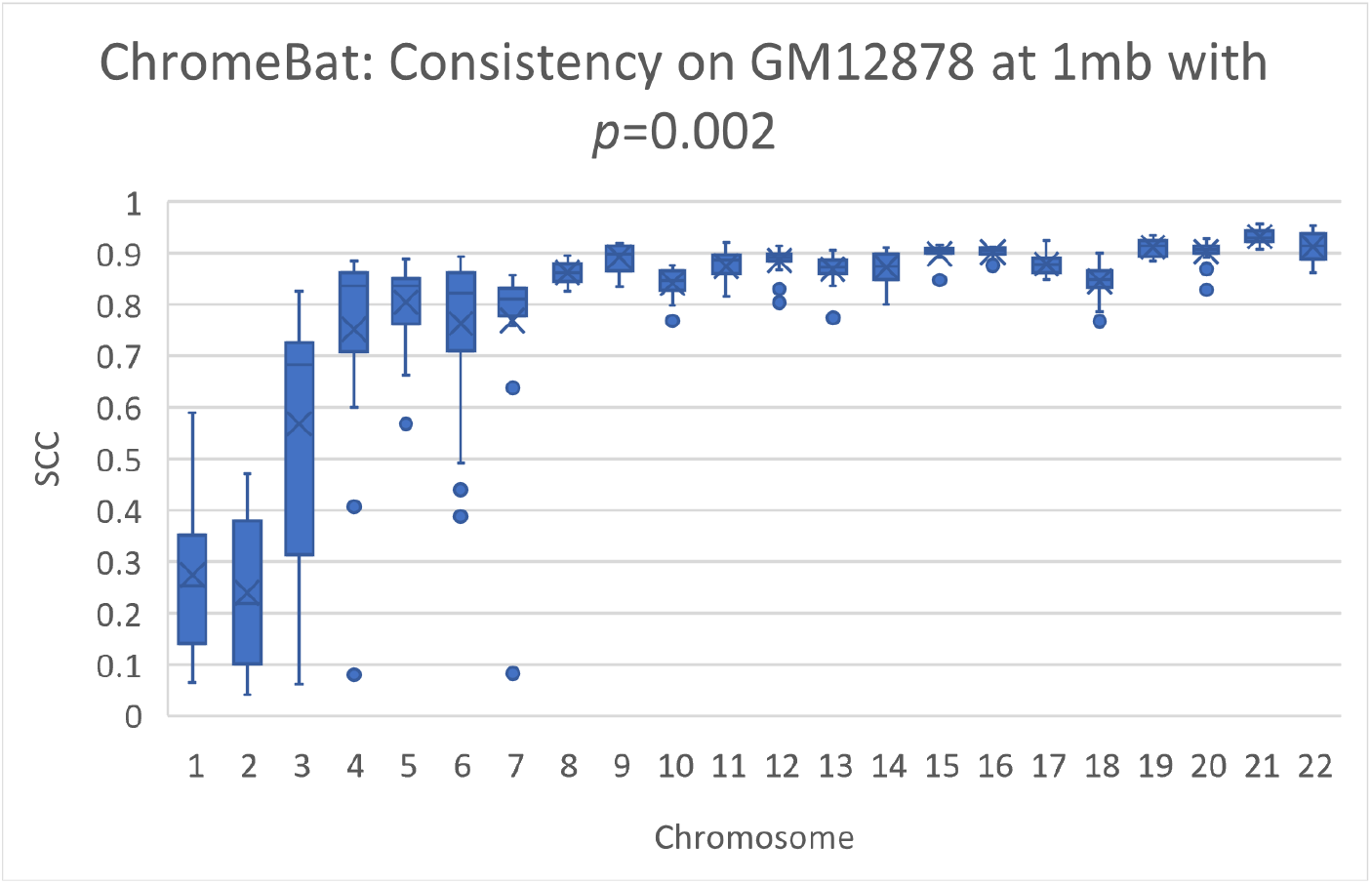
Consistency of ChromeBat on GM12878 with *p*=0.002. This is a consistency experiment done on GM12878 where hyperparameters from Table 1 with the exception of `structs=20` and *p* = 0.002.

**Figure 6.**
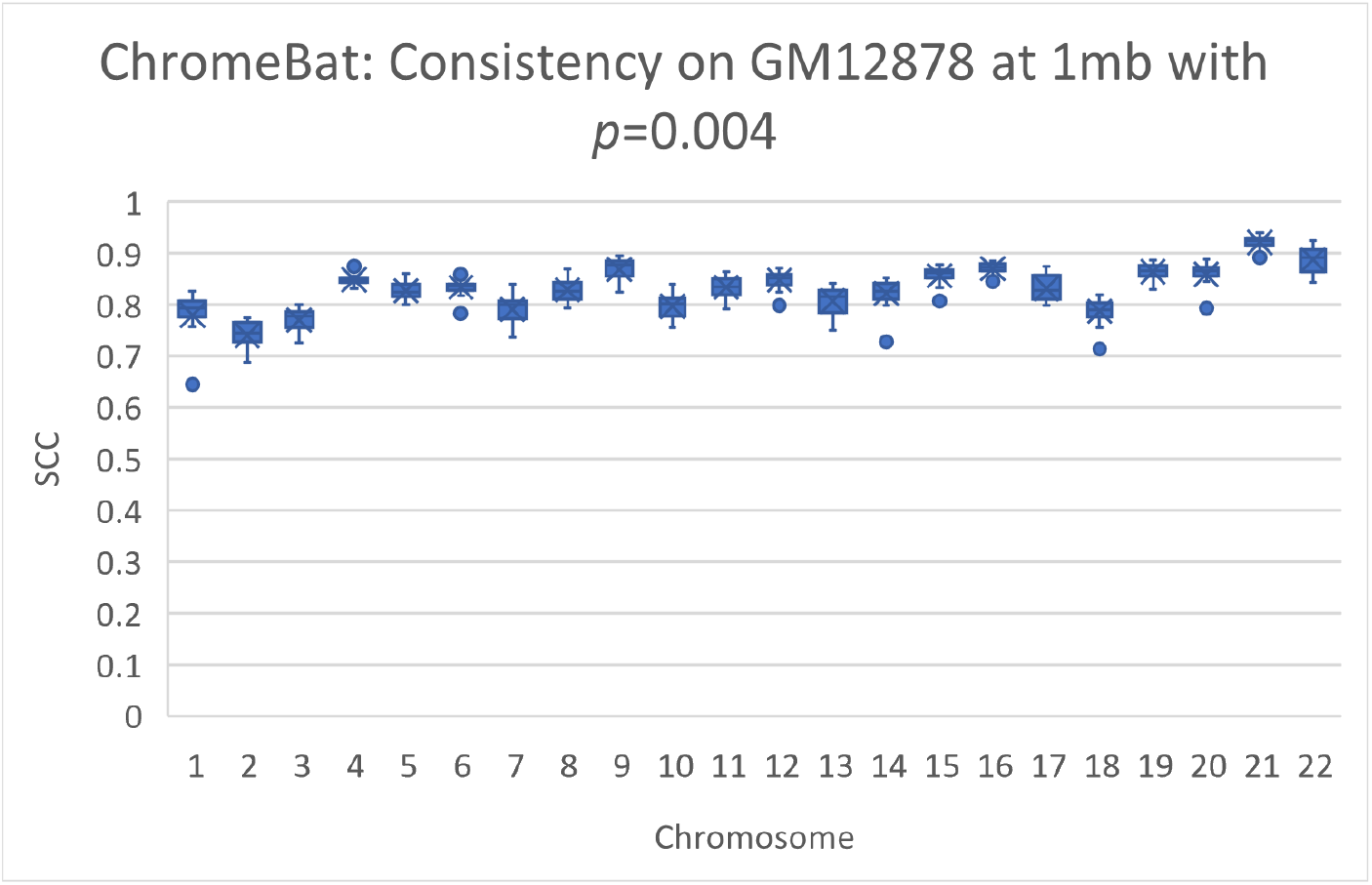
Consistency of ChromeBat on GM12878 with *p*=0.004.This is a consistency experiment done on GM12878 where hyperparameters from Table 1 with the exception of `structs=20` and *p* = 0.004.

### 2.4 Validation on FISH Data

We validate ChromeBat on FISH data. Chromosome 22 from GM06990 cell line was FISH probed at four loci in [5]. These loci (L5,L6,L7,L8) were gathered from consecutive positions in terms of base pairs but alternating between chromosome compartments A and B. In particular, (L5,L7) belongs to compartment A and (L6,L8) belongs to compartment B. Thus for our generated structure to be consistent with the FISH data we require (L5,L7) to be closer than (L5,L6) as well as (L6,L8) to be closer together then (L7,L8). As seen in Figure 7 we indeed have (L5,L7) to be closer than (L5,L6) as 2.2 *<* 2.7 and (L6,L8) is closer together then (L7,L8) as 2.3 *<* 2.9. This analysis and visualization was performed with PyMol [33].

**Figure 7.**
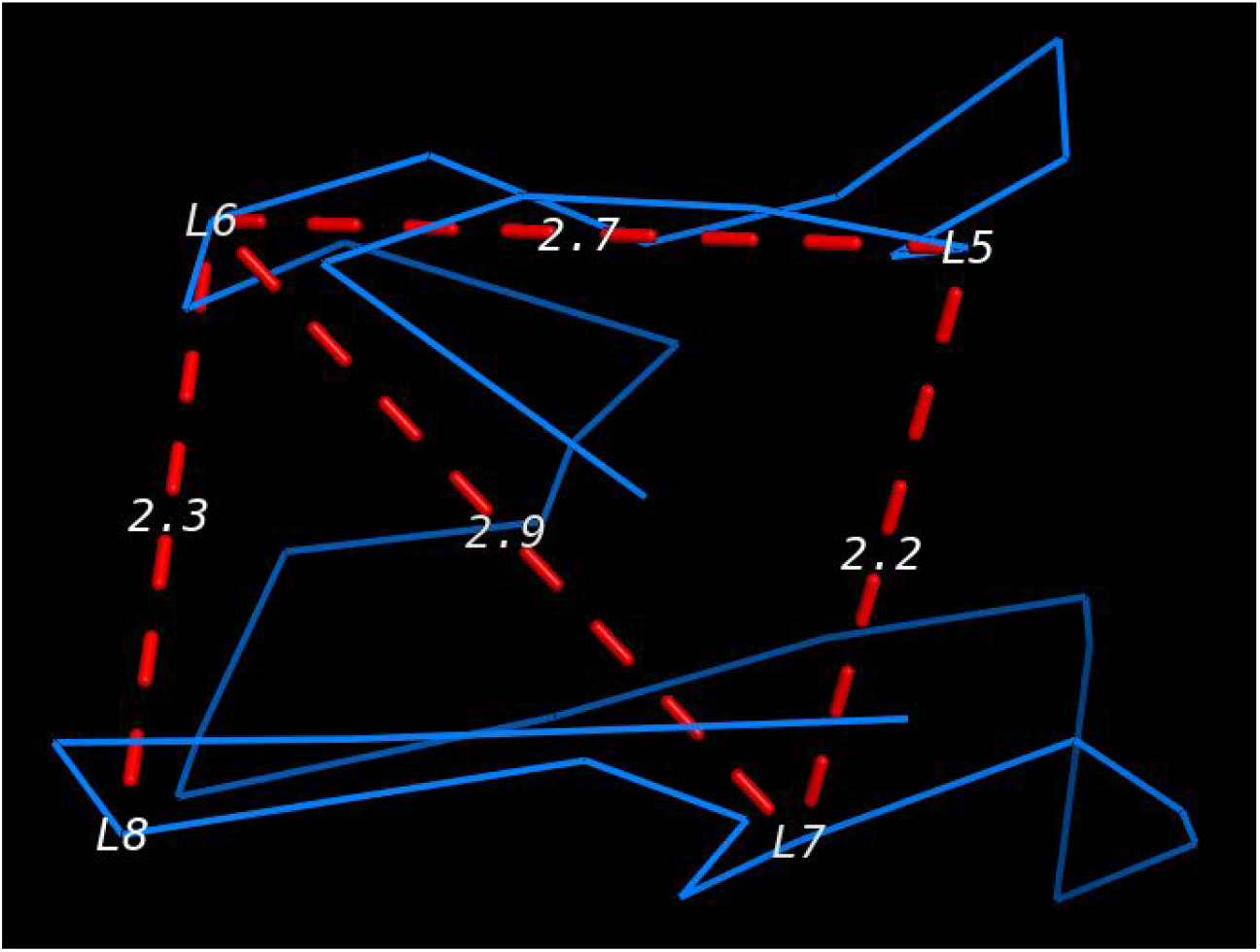
Verification of ChromeBat using FISH data. The blue line denotes chromosome 22 as reconstructed by ChromeBat on the GM06990 data. The four probes (L5,L6,L7,L8) from [5] are labeled on the chromosome. Additionally, the important distances between them have been labeled by red dashed lines.

## 3 Discussion

As highlighted in [9] the 3D-GRP lacks algorithmic diversity in general, however as ChromeBat is a metaheuristic approach, we restrict our attention to algorithmic diversity among metaheuristic algorithms. We found six metaheuristic methods in literature, however of those five are based on Simulated Annealing and one on the Genetic Algorithm. Figure 8 highlights this shortcoming categorically. It can be seen that these methods only represent two categories of metaheuristic algorithms, evolutionary and physics based. Thus, ChromeBat is the first representative of the nature-based methods, and many categories of metaheuristic algorithms are not studied on this problem.

**Figure 8.**
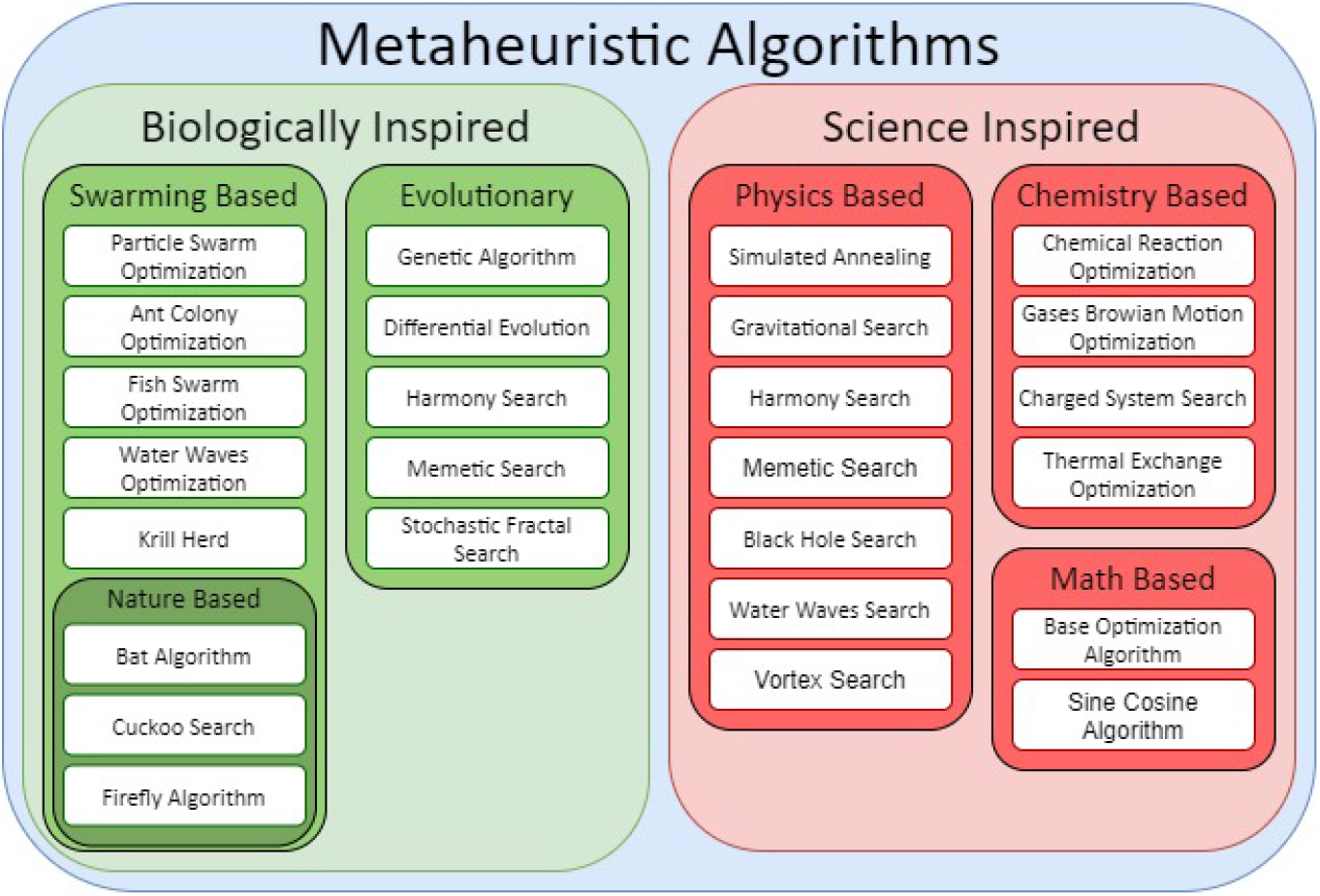
Classifications of Different Metaheuristic Algorithms. Compiled using information from [34, 35, 36, 37, 38, 39, 40].

Among metaheuristic algorithms the need for more algorithmic diversity in the 3D-GRP can be seen in Section 2.1. In particular, the comparison on GM12878 given in Figure 2 showcases that two different methods (HSA and ChromeBat) give state of the art performance on different chromosomes even within the cell line. These results, especially in combination with ChromeBat’s dominant performance on GM06990 seen in Figure 1, are empirical evidence that confirm the intuition that different methods have varying performance across different datasets. Compounded with how few metaheuristic methods are deployable on raw Hi-C data and poor characterization of what makes a method perform well on a given dataset, ChromeBat contributes diversity to a less-studied class of the 3D-GRP methods.

The importance of algorithmic diversity can be seen in a broader scale in Section 2.2. Particularly, on GM06990 shown in Figure 3 ChromeBat showcases competitive results across the board but of particular interest is its state of the art performance on chromosome 20. The fact that a bio-inspired approach performs well has two interpretations. First, the 3D-GRP domain might be best served using no singular method but instead an ensemble of methods for each task. This is due to the property that different methods appear to have different performances on different instances of the 3D-GRP, even in the same cell line. Secondly, it makes increasing the algorithmic diversity of studied methods more interesting as certain techniques may dominate portions of the 3D-GRP but no method will perform best across the entirety of the 3D-GRP. The proposed method seeks to advance the literature on both of these fronts.

### 3.1 Computation Time

We ran all presented results of ChromeBat using Intel(R) Xeon(R) CPU E7-4870 @ 2.40GHz with 1 Terabyte of RAM and 160 “processors” cores. On GM06990 the average computation time per chromosome was 773 seconds using the hyperparameters given in Table 1. In our implementation these hyperparameters are given in the `parameters_heavy.txt` file (available at https://github.com/OluwadareLab/ChromeBat). However, these parameters call for a search over 30 combinations of *α, p*, which in implementation becomes 30 concurrent processes. Because this could be computationally intense for most local machines, we also provide a `parameters_light.txt` file that reduces the searched *α, p* and will only open 6 concurrent processes.

## 4 Conclusion

We propose the development of ChromeBat Algorithm as a novel approach to solve the 3D-GRP. The domain in general lacks algorithmic diversity and so we base our approach in the bio-inspired Bat Algorithm. We find it is capable of state of the art performance on real Hi-C cell lines GM12878 and GM06990. This motivates future approaches to consider optimization algorithms that are metaheuristic in nature for the 3D-GRP domain and highlights interest in ensemble models that combine many approaches.

## 5 Methods

### 5.1 Loss Function

Let *A* ∈ **R**^*n*×*n*^ be a genome wide contact matrix obtained from a Hi-C experiment with *n* loci. We define genomic distance function between loci *i* and *j* as

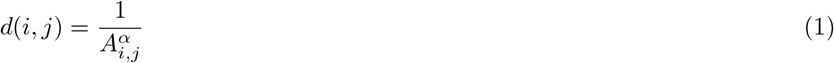

From Equation (1) we define distance matrix matrix *D*. That is, *D*_*i,j*_ = *d*(*i, j*) given contact matrix *A* and conversion factor *α* ∈ **R**.

Using *D*, we define the loss function as follows

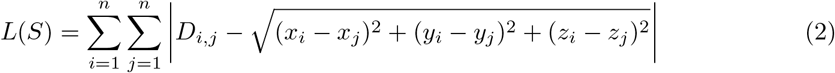

where *S* is a proposed structure with *n* (*x, y, z*) coordinates and *D*_*i,j*_ is defined. ChromeBat utilizes the loss function presented in Equation (2) that has been used in other works [1]. The loss function measures the difference between distance matrix *D* and the distance matrix induced by *S*.

### 5.2 Bat Algorithm For the 3D-GRP

Moving on to the Bat Algorithm (BA) for optimization, the algorithm is motivated as follows. Bats are largely blind predators that use echolocation to solve their objective of finding prey. They can alter their frequency and volume where high frequency yields a short range but high-resolution picture and vice versa for low frequency. They typically begin their search at high volume but then lower it as they draw near to their prey. Using these intuitions about how bats navigate the fundamental tradeoff of exploration versus exploitation, we present a summary of the Bat Algorithm, which is visualized in Figure 9. Note the following discussion describes how the BA is implemented in ChromeBat, which varies slightly from originally proposed. A list of hyperparameters for ChromeBat can be seen in Table 1. A description of the Bat Algorithm is as implemented in ChromeBat is as follows. Initialize *k* bats, where each bat *i* “knows” three locations, *S*_*i*_, 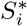, *S*\*, representing a bat’s current location, its personal best known location, and the global best location respectively. Note all of these vectors and velocity vector *V*_*i*_ are all in **R**^3*n*^, where *n* denotes the number of loci from the Hi-C experiment. We initialize 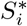 randomly, *V*_*i*_ as the the 0 vector, and *S*\* using Equation (7). The algorithm then proceeds for *T* iterations.

**Figure 9.**
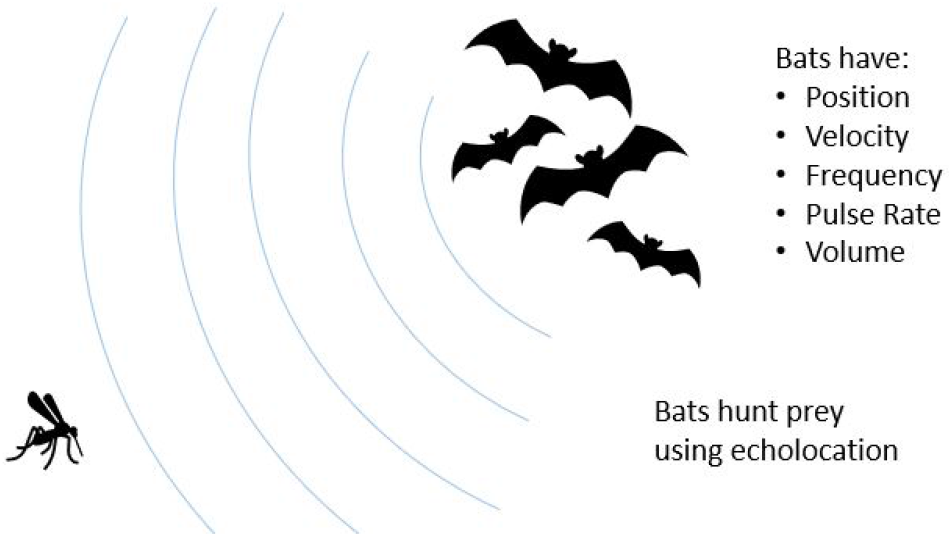
Visualization of the Bat Algorithm. The Bat Algorithm is inspired by the natural hunting behavior of bats. The algorithm captures this by giving each bat a the set of variables pictured on the right.

Each loop of the algorithm begins by the bats updating their current location *S*_*i*_ by the following equations:

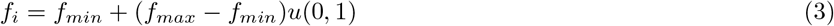

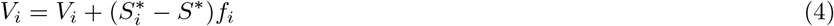

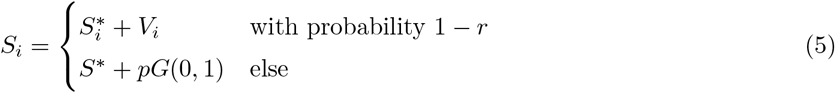

where *G*(*μ, σ*) denotes a vector of 3*n* values where each one is sampled from a normal distribution with mean *μ* and standard deviation *σ*, and *u*(*a, b*) denotes a value selected uniformly at random from the interval [*a, b*]. Note by the random valued condition in Equation (5) bats update their position in one of two ways. If they decide not to pulse (corresponding to probability 1 − *r*), then they will select a random frequency in [*f*_*min*_, *f*_*max*_] as per Equation (3), and use this value to randomly adjust their velocity *V*_*i*_ in Equation (4). This velocity adjustment is based on a bats current best location 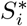 and global best location *S*\*, guiding bats in a hopefully well selected direction. Then the bats who did not pulse will use their newly updated velocity to update their current position *S*_*i*_. On the other hand if a bat chooses to pulse, invoking the second case in Equation (5), then the bat will teleport to the global best known location *S*\* and take a random walk scaled by hyperparameter *p*. It can be seen that a high *r* corresponds to a bat who pulses with high probability and vice versa for low *r*.

Once bats have updated their location *S*_*i*_ they decide whether or not to accept their new solutions to the 3D-GRP according to equations:

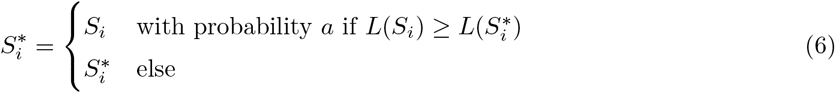

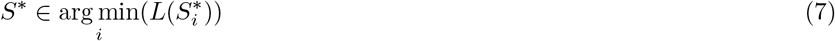

where *L* is the loss function defined in Equation (2). The conditions on whether a bat accepts its current solution *S*_*i*_ are given in the top case of Equation (6) and can be interpreted as follows. For a solution *S*_*i*_ to accepted, it must be a better solution then the current accepted solution solution 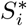, that is 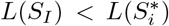. Additionally, the bat’s volume *a* as is used to introduce randomness in whether a bat accepts solutions frequently. From Equation (6) it can be seen that a high *a* means a bat will accept new solutions with high probability. The final step of the algorithm is to reapply Equation (7) to select the new global best solution *S*\*. After all *T* iterations are complete the algorithm terminates and returns *S*\*.

In general, there is a tradeoff in non-convex optimization between exploration and exploitation. The success of this algorithm comes from its balancing act between the two. The algorithm has been successfully applied in many problem domains such as continuous optimization, parameter estimation and image processing [3], and so we expect it to perform well on the 3D-GRP. As mentioned earlier, the 3D-GRP is a high dimensional problem, so we will also apply variants of the Bat Algorithm suited for this context.

### 5.3 Hyperparameter Selection

As seen in Table 1 ChromeBat features many hyperparameters. To select default values for the algorithm we conduct a series of experiments on the simulated helical structure data presented in [17]. More details on this data set can also be found in Section 5.5, and all experiments are performed on the 90% coverage version.

Initially, we take *f*_*min*_ = 0 as it is intuitive by Equation (3) that bats should select frequency uniformly between 0 and *f*_*max*_. Then we perform four searches across with *f*_*max*_, *p*, *r*, and *a* at the same time. The details about how these parameters were assigned through searches can be found in the provided additional files [ see Additional file 1, Additional file 2, Additional file 3]. We find in all of these searches that *α* = 0.5 so we may fix it for future searches. Additionally, we find the greatest Spearman Correlation Coefficient (SCC) result in occurred at *p* = 0.9 and *a* = 0.9 so we fix these parameters [see Additional file 3].

With *α* known we conduct an experiment searching across *T* and *k* as these parameters solely determine the runtime of the algorithm. In the Additional file 4, we find *k* = 10 and *T* = 10, 000 is sufficient, noting the differences in SCC between runs with greater *T, k* is negligible. We also notice from this experiment that no matter large *T* and *k* the algorithm appears to get “stuck” sometimes. To remedy this we introduce another hyperparameter `structs`, that represents how many structures the algorithm should generate for consistency. We take `structs = 10` to balance computation time and performance of the algorithm. Further discussion of this parameter can be seen in Section 2.3.

Finally, we carry out a search across *f*_*max*_, *p* that can be seen in the Additional file 5, where we take generate 10 structures per parameter combinations due to concerns about the methods consistency from the previous search. To ensure consistency we use the average SCC across the 10 generated structures and the most consistent and best performance from *f*_*max*_ = 0.1 and *p* = 0.002. Thus, we fix *f*_*max*_ = 0.1 but we find impressive performance across all *p* values searched {0.002, 0.004, 0.006, 0.008, 0.01}. Thus for the default behavior of the method we include a search across *α* and *p*.

### 5.4 Evaluation

To validate our method we use the Spearman Correlation Coefficient metric. The equation for this metric is given by:

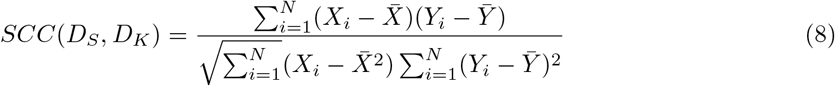

Where *D*_*K*_ is a set containing all unique distance measures between loci from the Hi-C experiment. That is, *D*_*K*_ has an element for each unique defined entry of *D*. Let *D*_*S*_ be a set of all corresponding distance measures, particularly that *D*_*S*_ contains the distance measure between loci *i, j* only if *D*_*i,j*_ is defined. Then let *X, Y* be ranked variables corresponding to *D*_*S*_, *D*_*K*_ respectively and let 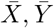 refer to the mean of the ranked variables.

### 5.5 Datasets

To demonstrate the effectiveness of ChromeBat, we compare the method with techniques studied in the literature on two cell lines and a simulated dataset. The first cell line is the normalized GM12878 cell line [41] downloaded from GSDB [42]. The normalization method used is the Knight-Ruiz [43]. This cell line contains 4.9 billion pairwise contacts at map resolution 950 bp. It was gathered from Human GM12878 B-lymphoblastoid cells, aggregated from 9 cultures. The second cell line is GM06990 normalized with Yaffe and Tanay [44]. The GM06990 cell line was litigated with Hind III enzyme and downloaded from http://sysbio.rnet.missouri.edu/bdm_download/3DEM/. Both cell lines are considered at 1mb resolution.

In addition to assessing ChromeBat’s performance on real data, we tune its hyper-parameters using simulated data from [17]. This data was constructed by simulating a regular helical structure and deriving contacts maps at a desired signal coverage level. A signal coverage level merely denotes what percentage of entries in the contact matrix are non-zero. Zhang et al [17] derives contact matrices at a desired signal coverage level by assuming the contact matrix satisfies a Poisson distribution of a power law based on the actual distances. Then they provide and test their methods on coverage levels of 90%, 70%, and 25%. The motivation for the simulated approach is that genome-scale ground truth exists for any genome reconstruction problem. Because of this, we use the simulated data set for hyperparameter selection of our model.

## Appendix

Not Applicable

## Abbreviations

bp: base pairs
3D: Three Dimensional
NGS: Next Generation Sequencing
TAD: Topologically Associating Domain
3D-GRP: Three Dimensional Genome Reconstruction, Problem
MDS: Multidimensional Scaling
LAD: Lamina-Associated Domain
SCC: Spearman Correlation Coefficient
SA: Simulated Annealing
PSO: Particle Swarm Optimization

## Acknowledgements

Not Applicable

## Availability of data and materials

The GM06990 cell line analysed during the current study are available here, http://sysbio.rnet.missouri.edu/bdm_download/3DEM/3DMax_inputs/GM06990_SquareMatrix_Format/GM06990_Yaffe&Tanay_normalized/HindIII/.

The GM12878 cell line analysed during the current study are available in the GSDB repository, http://sysbio.rnet.missouri.edu/3dgenome/GSDB/details.php?id=GM12878.

The simulated data set analysed during the current study are available in the Ouyang Lab repository, https://people.umass.edu/ouyanglab/hsa/downloads.html#Data.

The ChromeBat method generated and analysed during the current study are available in the Oluwadare Lab repository, https://github.com/OluwadareLab/ChromeBat.

## Ethics approval and consent to participate

Not Applicable

## Consent for publication

Not Applicable

## Authors’ contributions

O.O. conceived the project. O.O. and B.C. developed the method. B.C. programmed the method and wrote the manuscript. O.O. and P.B. reviewed and revised the manuscript. All authors read and approved the final manuscript.

## Author details

Department of Computer Science, University of Colorado Colorado Springs, Colorado Springs, USA.

## Additional Files

Additional file 1 — Frequency Search

The file is a spreadsheet in .xlsx format that details how the frequency *fmax* parameter was determined through a search of conversion factor *α* versus frequency *fmax* on the simulated Hi-C data.

Additional file 2 — Perturbation Search

The file is a spreadsheet in .xlsx format that details how the perturbation *p* parameter was determined through a search of conversion factor *α* versus perturbation *p* on the simulated Hi-C data.

Additional file 3 — Pulse and Volume Search

The file is a spreadsheet in .xlsx format that details how the pulse *r* and Volume *a* parameter were determined through a search of conversion factor *α* versus pulse *r* and the volume *a* parameters on the simulated Hi-C data.

Additional file 4 — Number of Bats versus Number of Iterations Search

The file is a spreadsheet in .xlsx format that details how the number of bats *k* and the number of iterations *T* through a search of bats *k* versus iterations *T* parameters on the simulated Hi-C data.

Additional file 5 — Frequency versus Perturbation Search

The file is a spreadsheet in .xlsx format that details the effect of changes in frequency *fmax* and perturbation *p* on the performance of our algorithm on the simulated Hi-C data

Additional file 6 — Spearman Correlation of all Methods

The file is a spreadsheet in .xlsx format that reports the SCC of all compared methods on all chromosomes of GM06990.

Additional file 7 — Spearman Correlation of all Methods

The file is a spreadsheet in .xlsx format that reports the SCC of all compared methods on all chromosomes of GM12878.

